# SOX9 elongates cell cycle phases and biases fate decisions in human intestinal stem cells

**DOI:** 10.1101/2022.11.03.514885

**Authors:** Joseph Burclaff, R. Jarrett Bliton, Keith A Breau, Michael J Cotton, Caroline M Hinesley, Meryem T Ok, Caden W Sweet, Anna Zheng, Eric D Bankaitis, Pablo Ariel, Scott T Magness

**Affiliations:** Joint Department of Biomedical Engineering, University of North Carolina at Chapel Hill/North Carolina State University, Chapel Hill, North Carolina; Center for Gastrointestinal Biology and Disease, University of North Carolina at Chapel Hill, Chapel Hill, North Carolina; Department of Cell Biology and Physiology, University of North Carolina at Chapel Hill, Chapel Hill, North Carolina; Microscopy Services Laboratory, Department of Pathology and Laboratory Medicine, University of North Carolina at Chapel Hill, Chapel Hill, North Carolina; Lineberger Comprehensive Cancer Center, University of North Carolina at Chapel Hill, Chapel Hill, North Carolina; Department of Genetics, University of North Carolina at Chapel Hill, Chapel Hill, North Carolina

## Abstract

**Background and Aims:** The transcription factor SOX9 is expressed in many stem/progenitor cell populations and has biphasic correlations with proliferation rates across different biological systems. In murine intestinal crypts, distinct Sox9 levels mark three phenotypically different cell types, with lowest levels marking rapidly-dividing transit amplifying (TA) cells, intermediate levels marking intestinal stem cells (ISCs), and highest levels marking slowly-dividing label retaining secretory precursors. SOX9 expression levels and the impact of these levels on cell cycle and stem cell activity have not been characterized for humans.

**Methods:** Monolayers of primary human ISCs isolated from healthy organ donors were engineered with stable SOX9-knockout (KO) and/or SOX9-overexpression (OE) genomic modifications to assess the impact of SOX9 levels on proliferative capacity by DNA content analysis, cell cycle phase length by live imaging for a PIP-FUCCI reporter, stem cell activity via organoid formation assays, and cell fate after ISC differentiation tracked via qPCR.

**Results:** SOX9 was expressed at diverse levels in human intestinal crypt lineages in vivo, repressed proliferation in human ISC monolayers, and predominantly lengthened G1 by >40% with lesser lengthening of S and G2/M phases. Overexpression of SOX9 caused slower proliferation yet increased organoid forming efficiency. Higher SOX9 levels biased ISC differentiation towards tuft cell and follicle-associated epithelium fates while loss of SOX9 biased cells toward absorptive enterocyte, goblet cell, BEST4^+^ cell, and enteroendocrine cell fates.

**Conclusions:** SOX9 is a master regulator of stem cell activity in human ISCs, lengthening the cell cycle, promoting stemness, and altering differentiation fate. Interestingly, differences are noted between species, highlighting the importance of analyzing regulatory mechanisms in primary healthy human cells.

## INTRODUCTION

The small intestinal crypt represents one of the most proliferative tissues in mice and humans, with complete turnover of the intestinal epithelium suggested to occur every 5-7 days. This rapid epithelial self-renewal is facilitated by distinct proliferative epithelial populations within the crypt including the active intestinal stem cell (ISC) at the crypt base, transit amplifying (TA) cells, and putative quiescent reserve ISCs^1^. These cell populations have different proliferative capacities, with active ISCs proliferating about once daily, TA cells cycling multiple times per day as they rise from the crypt and differentiate^1^, and putative reserve ISCs being largely quiescent at homeostasis yet re-initiating proliferation to repopulate the crypt upon injury to active ISCs^2-11^.

The decision between ISC self-renewal (remaining in cell cycle) and differentiation (leaving cell cycle) is inherently linked to cell identity and fate, but mechanisms underpinning this critical decision are not clear. Central to elucidating these mechanisms is determining how and when the distinct lengths of G1, S, and G2/M cell cycle phases influence ISC cell fate decisions. Our current understanding of mammalian ISC properties and function comes almost exclusively from murine models. However, increasing studies are finding that murine and human ISCs exhibit fundamental differences at transcriptomic and functional levels^12-14^. This raises questions surrounding the physiological equivalence of self-renewal and cell fate decisions between murine and human ISCs and provides merit to evaluate ISC maintenance and differentiation mechanisms in human ISCs.

SRY-Box Transcription Factor 9 (Sox9) is known to regulate stem cell maintenance and differentiation in many adult tissues in mice including the intestine, colon, pancreas, and liver^15-17^. Previous work from our lab described distinct expression levels of Sox9 present across proliferative and dormant cell populations within the murine small intestinal crypt^7, 18-20^. Using a Sox9-EGFP reporter mouse model and immunostaining for endogenous SOX9, we demonstrated that cells with high levels of Sox9 (Sox9-high) within the crypt are enriched for enteroendocrine cell markers and are consistent with label retaining cells (LRCs)^21^ shown to have reserve ISC (rISC) function^19, 20^. Sox9-low cells represent actively proliferating ISCs (aISCs), and even lower Sox9 levels termed Sox9-sub-low mark rapidly dividing TA cells^18^. Sox9 is undetectable in enterocyte and goblet cell lineages, but high Sox9 levels are expressed in 50% of Paneth cells in the crypt base and 100% of tuft cells in the villus^19^, highlighting a role for high levels of Sox9 in select differentiated cell types.

In mice, genetic ablation of Sox9 results in substantially reduced numbers of Paneth cells and a less appreciable reduction in enteroendocrine cells, further supporting a role for Sox9 in lineage allocation^22, 23^. Related to this finding, our prior work demonstrates that high levels of Sox9 are expressed in LRCs shown to be secretory progenitor cells that predominantly give rise to Paneth cells and some EECs in homeostasis^21^. After radiation damage, these LRCs have potential to revert to an active ISC state, underscoring the plasticity within this cell population^21^. Other work by our group demonstrated that cells expressing high Sox9 levels in the crypt typically do not divide in mice, but subsets of Sox9-high cells co-express Ki67 after radiation during the crypt regeneration phase, indicating that they retain proliferative capacity under injury conditions and may support re-entry into an active cycling state^20^. Together, these observations raise the question as to whether differential Sox9 levels regulate cell cycle phase lengths across these cell types, and moreover, whether a similar correlation exists between SOX9 levels, proliferation rates, and cell fate decisions in the equivalent human cell populations.

Although mechanisms focused on cyclins, cyclin-dependent kinases, and retinoblastoma (Rb) have a firm foundation in defining key decision points for cycling cells to enter subsequent phases, upstream transcriptional influences on phase length and progression are less understood. This is especially the case regarding how upstream transcription factors modulate important restriction points in the cell cycle that ultimately influence cell cycle exit, differentiation, and maturation. For instance, in the developing mouse retina, a graded reduction of Sox2 levels causes premature cell cycle exit of retinal progenitors, resulting in aberrant neural progenitor differentiation that is directly proportional to the Sox2 levels^24^. This example underscores that transcription factor levels can be crucial for spatial and temporal cell cycle decisions that regulate cell differentiation, maturity, and ultimately function.

In particular, the G1 phase of the cell cycle seems to be a permissive state for cell fate decisions and perhaps maturation^25, 26^. Several studies show that embryonic stem cells re responsive to differentiation cues during the G1 phase^27-29^, highlighting this as an important window of time whose length may be critical to fully specify cell lineages or keep cells in a more permissive and plastic state. Here, we analyze how SOX9 regulates human ISC cell cycle activity and fate decisions. We design a genetic and cellular toolkit allowing complete control of SOX9 expression in cultured primary human ISCs, then analyze its effect on proliferation, cell cycle, stemness, and differentiation.

## RESULTS

### SOX9 is expressed at different levels in human intestinal crypt lineages

Since all published reports detailing expression levels of SOX9 within the intestinal crypt lineages have come from mice^7, 18-20^, we stained for SOX9 expression in the human intestinal crypt (Fig 1A). Similar to the mouse crypt, differing levels of SOX9 were seen across cell types of the human crypt, with moderate levels of SOX9 in cells at the crypt base, low yet detectable levels throughout the KI67^+^ TA zone, and SOX9-high Paneth cells at the crypt base. No work has defined correlates to the murine SOX9-high LRCs in humans, though the high SOX9 expression in Paneth cells suggests they or their precursors may share attributes with this population. Notably, all proliferating populations expressed detectable SOX9 protein, consistent with the mouse intestine. To validate this trend in three additional human samples, we checked *SOX9* expression in small intestinal ISCs and TA cells from our recently published database of single cell RNA sequencing (scRNAseq) results from different regions of the intestine across three healthy human donors^13^. As expected, we found that ISC markers *LGR5* and *OLFM4* marked ISCs, proliferation markers *MKI67* and *PCNA* were enriched in TA cells, and *SOX9* had higher expression in ISCs, with lower but still notable expression in TA cells (Fig 1B). This supports that SOX9 marks proliferative human crypt populations consistent with what has been experimentally described in mouse intestine.

**Figure 1:**
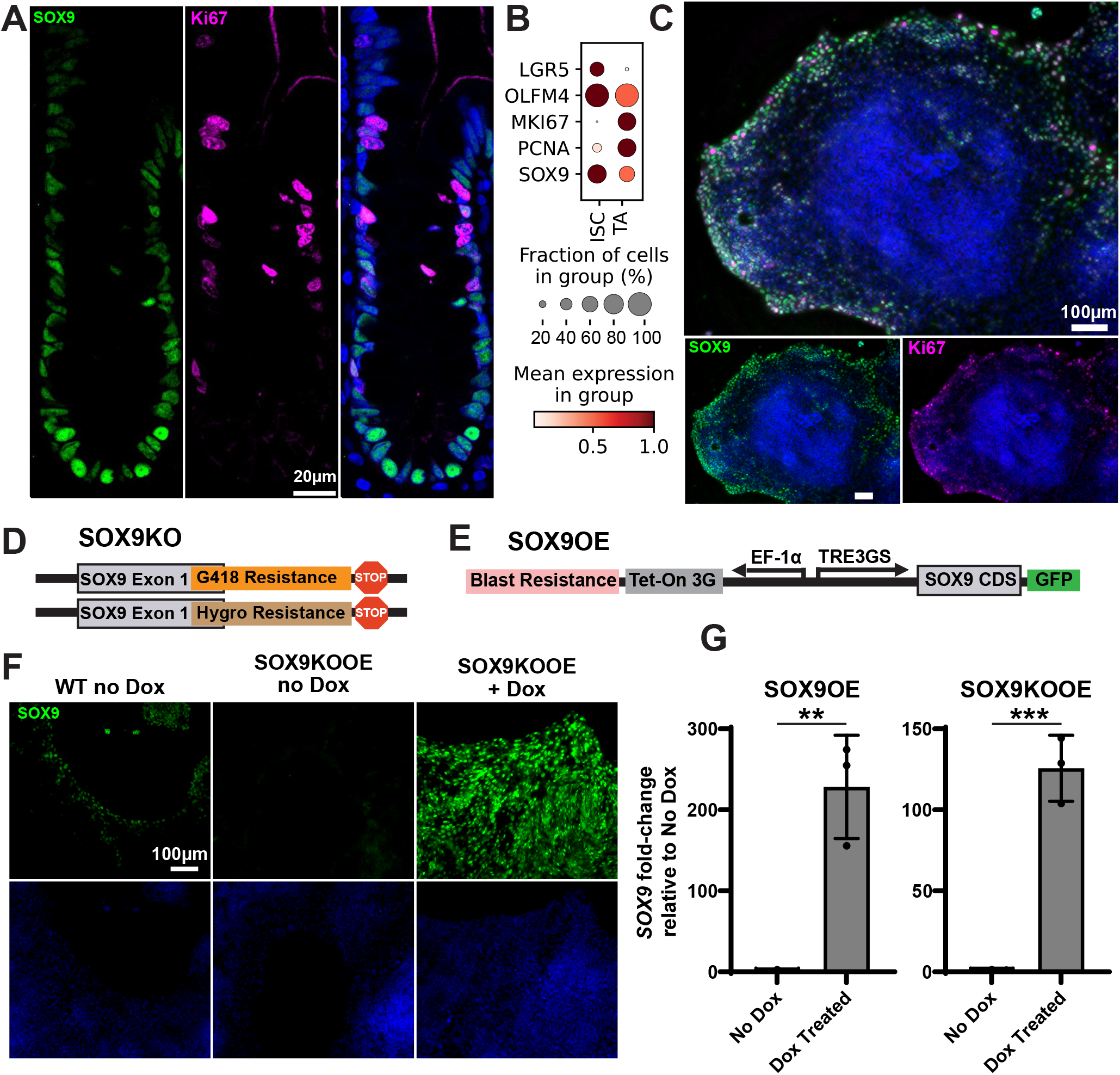
SOX9 expression and modulation within human ISCs. **A)** Immunofluorescence staining for SOX9 (green) and KI67 (pink) in a human jejunal crypt. Scale bar = 20 μm. **B)** Expression of markers of stemness (*LGR5, OLFM4*), proliferation (*MKI67, PCNA*) and *SOX9* across intestinal stem cell (ISC) and transit amplifying (TA) cells of the small intestine using scRNAseq data from three healthy human donors^13^. **C)** Immunofluorescence staining for SOX9 (green) and KI67 (pink) in an expanding monolayer of human jejunal stem cells. All scale bars = 100 μm. **D)** Diagram of SOX9-knockout (KO) constructs, using CRISPR/Cas9 to insert two selectable antibiotic resistance genes followed by stop codons into exon 1 of SOX9. **E)** Diagram of SOX9-overexpression (OE) construct, using PiggyBac transposase to insert a Tet-On system for driving doxycycline (Dox)-inducible *SOX9* expression into the genome. **F)** Immunofluorescence staining for SOX9 (green) in wildtype (WT), SOX9KOOE no Dox, and SOX9KOOE+Dox monolayers. Scale bar = 100 μm. **G)** qPCR results showing relative expression of *SOX9* in SOX9OE no Dox and SOX9OE+Dox monolayers (left) or SOX9KOOE no Dox and SOX9KOOE+Dox monolayers (right), with all values normalized to expression levels without Dox. All cells were grown in differentiation media with/without Dox for 5 days. Statistics shown using T Test, P values: * < 0.05, ** <0.005, *** < 0.001.

### A 2D monolayer system enables ISC culture and supports generation of human ISCs with tunable SOX9

To study the effects of SOX9 in human ISCs, we cultured primary jejunal crypts from a recently deceased human organ donor on collagen patties to grow monolayers as described in our prior work^30-32^. As a first pass to show that SOX9 is expressed similarly in this in vitro environment as it is in the crypt, we stained for SOX9 and KI67 in actively growing human monolayers cultured in Maintenance Media (MM)^30^ (Fig 1C). Monolayers grown in this media proliferate at the expanding edges, with little proliferation in the central cells. SOX9 was expressed across the proliferative region, with SOX9 present in all KI67^+^ proliferating cells. These findings show that the 2D monolayer conditions replicate the physiological crypt in that SOX9 is expressed in all proliferative populations of the growing intestinal epithelium.

Using our recently published optimized electroporation protocols for primary human intestinal monolayers^33^, we created transgenic ISC lines as a genetic toolkit for assaying how different levels of SOX9 affect human ISC activity. For a SOX9-null ISC line, CRISPR/Cas9 was used to knock out both SOX9 alleles with dual selectable resistance genes (SOX9KO, Fig 1D). For an inducible SOX9 ISC line, the PiggyBac Transposase system was used to stably introduce a doxycycline (Dox)-inducible SOX9 overexpression cassette into the genome (SOX9OE, Fig 1E). Stable lines were made for both SOX9KO and SOX9OE independently, and we created a third SOX9 rescue line (SOX9KOOE) where the SOX9OE construct was stably integrated into the SOX9KO line. Control of tunable SOX9 expression was validated via immunofluorescence (Fig 1F) and qPCR (Fig 1G). Control ISCs demonstrated robust SOX9 staining at the leading edges of the colonies as expected. No specific SOX9 staining was observed in SOX9KO ISCs, with known background staining observed on colony edges^34^, and SOX9OE ISCs demonstrated high levels of SOX9 across all cells in the ISC colony (Fig 1F). Consistent with the immunostaining, qPCR demonstrated proper *SOX9* overexpression, with >100x more *SOX9* transcript observed after treating differentiated cells with 100 ng/mL Dox. These ISC lines present the opportunity to study the effects of SOX9 on ISC activity across a wide range of SOX9 expression levels.

### SOX9 represses proliferation in human ISCs

We first explored how a range of SOX9 levels would impact proliferation by evaluating the number of cells in S phase after a single one-hour EdU pulse. Flow cytometry demonstrated 45.3% more SOX9KO ISCs took up EdU than WT ISCs, suggesting more SOX9KO cells were actively proliferating (Fig 2A). The inverse was seen with high SOX9, with 44.5% and 23.9% less cells taking up EdU in SOX9OE and SOX9KOOE cells, respectively, following Dox treatment (Fig 2B). Importantly, Dox had no discernable effect on WT or KO cells (Fig 2B), indicating that the proliferation changes are a direct consequence of the altered SOX9 levels.

**Figure 2:**
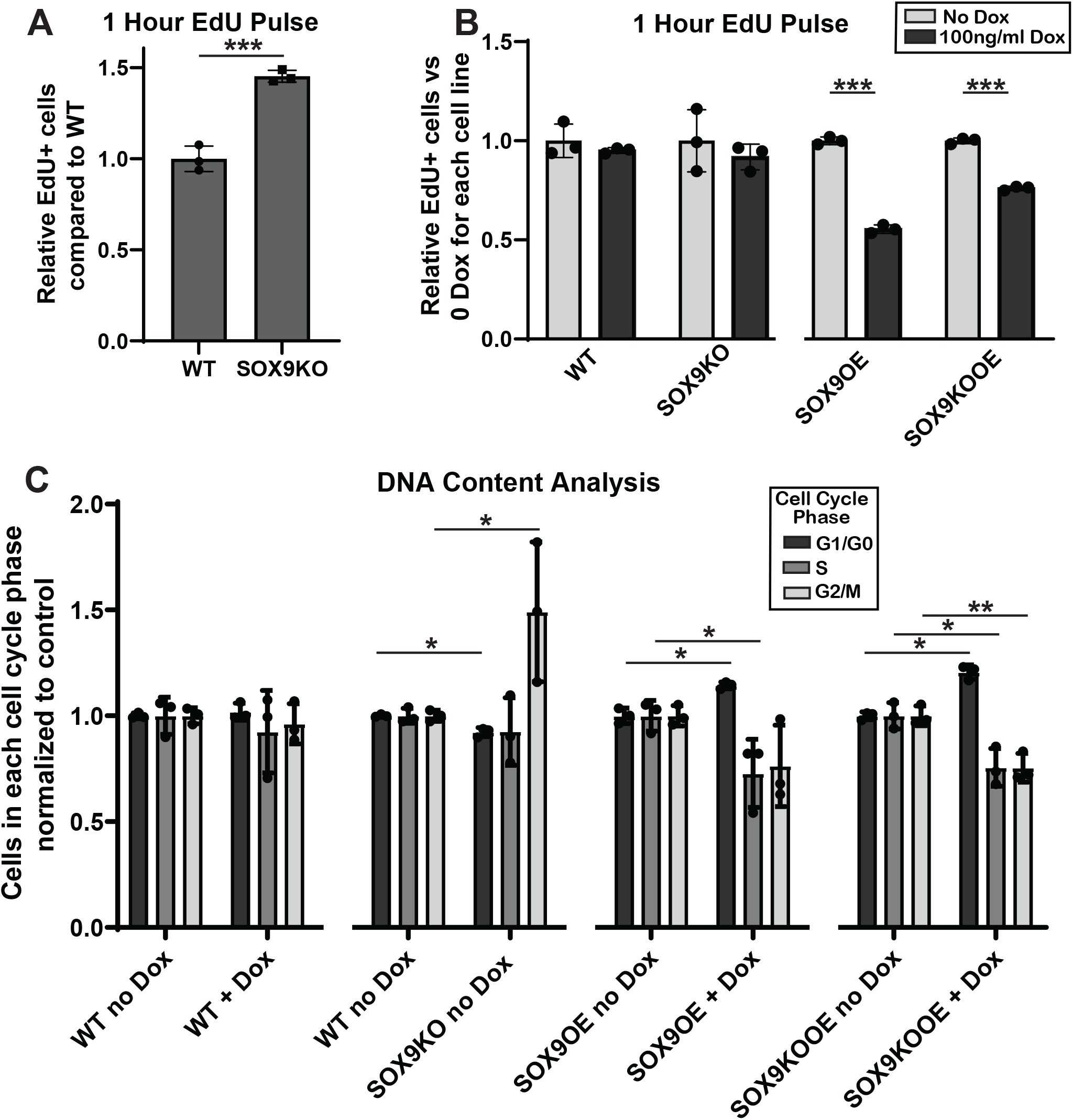
SOX9 controls proliferation and cell cycle in human ISCs. **A)** Number of cells positive for EdU staining following a 1 h EdU pulse in WT and SOX9KO monolayers. Values quantified via FACS and shown relative to WT levels. **B)** Number of cells positive for fluorescence staining for EdU following a 1 hr EdU pulse in WT, SOX9KO, Sox9OE, and Sox9KOOE monolayers all without dox (left bar of each pair, light gray) and with Dox added (right bar, dark gray). Values quantified via FACS and all values shown relative to the no Dox control for each cell line to demonstrate proliferation differences induced by Dox-driven SOX9. **C)** DNA content analysis using propidium iodide to track cell cycle fate within individual cells from WT, SOX9KO, SOX9OE, and SOX9KOOE cells all with and without Dox. Values were quantified via FACS. Bar charts represent proportion of cells calculated to be within each cell cycle phase: G1 (left), S (middle), G2/M (right) relative to the control for each condition. Statistics shown using T Test, P values: * < 0.05, ** <0.005, *** < 0.001.

To probe how SOX9 expression levels affect the distribution of cells across cell cycle phases, we treated growing monolayers with/without Dox for 48h then dissociated, fixed, and performed flow cytometry for total DNA content using Propidium Iodide (PI) (Fig 2C). To minimize differences due to contact inhibition, all monolayers for direct comparisons were plated side-by-side with similar cell seeding densities. We first looked at WT cells +/-Dox and found that 100 ng/mL Dox had no discernable effect on cell cycle. Consistent with their higher proliferation rate (Fig 2A), less SOX9KO cells were in G1/G0 phase, with an increase in G2/M phase cells and no statistical difference in S phase (Fig 2C). SOX9OE cells with Dox-driven SOX9 expression showed the opposite effect, with more cells in the G1/G0 phase and less in S phase compared controls. SOX9KOOE cells were affected in a manner consistent with SOX9OE, with more cells in G1/G0 and fewer in G2/M and S phases following Dox-induced SOX9 expression. All three conditions are consistent with a model where higher SOX9 levels lengthen total cell cycle largely by increasing G1 phase length.

### SOX9 elongates all cell cycle phases in human ISCs with preferential lengthening of G1

A major caveat of flow cytometry for DNA content is that it does not directly measure lengths of cell cycle phases. Also, since cells in the quiescent G0 phase and those in the proliferative G1 phase have the same DNA content, they cannot be distinguished using DNA content analysis. For example, a smaller proportion of cells in G1/G0 phase may indicate a shorter G1 phase in actively cycling cells and/or more cells actively cycling (less in G0) but no change in G1 length in the cycling cells. To quantify how SOX9 controls the length of cell cycle phases in actively cycling human ISCs, we used the recently published PIP-FUCCI allele as a fluorescent readout for all phases of the cell cycle^35^. This reporter consists of two alleles: a Cdt1_30-120_-mVenus fusion that has strong fluorescence in G1, rapid degradation at the start of S Phase, then increasing fluorescence in G2 phase through cytokinesis; and a Gem_1-110_-mCherry fusion with no fluorescence in G1, increasing fluorescence throughout S phase, then high fluorescence in G2/M until cytokinesis, at which point it rapidly dims (Fig 3A). This reporter was chosen for its distinct changes before and after G1 allowing for accurate quantification of G1 length using live-cell imaging.

**Figure 3:**
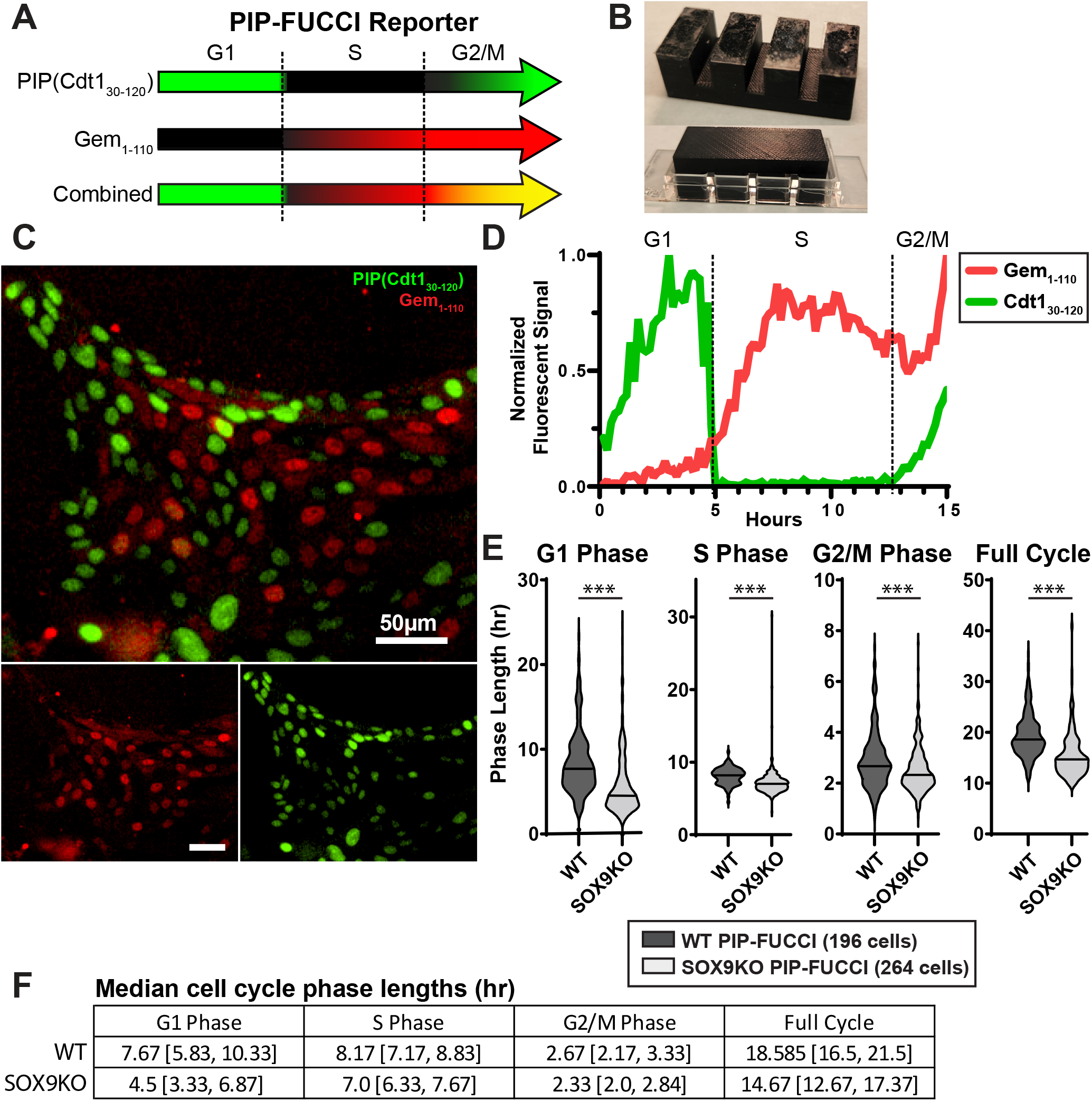
Quantifying cell cycle phase lengths. **A)** Schematic of dynamic reporter colors throughout the cell cycle in PIP-FUCCI cells. **B)** 3D printed collagen press used to make flat collagen layers in chamber slides for optimal live imaging on an inverted confocal microscope. **C)** Representative fluorescence image of the PIP-FUCCI reporter in expanding human jejunal monolayers. All scale bars = 50μm. **D)** Example fluorescence data resulting from tracking an individual PIP-FUCCI nucleus over its 15-hour cell cycle. **E)** Violin plots denoting lengths of each cell cycle phase across 196 WT and 264 SOX9KO cells as quantified using the PIP-FUCCI reporter. Median values are denoted on each violin plot. **F)** Median values with [Q1, Q3] quartile values are shown for each cell cycle phase and total cell cycle for WT and SOX9KO cells.

We engineered the PIP-FUCCI allele onto a PiggyBac transposase plasmid, transfected this plasmid into WT or SOX9KO cells^33^, and selected stable clones. As ISC monolayers necessitate a thick layer of soft collagen to achieve the ∼100 Pa stiffness needed to maintain stemness and monolayer spreading^31, 32^, we coated a cover-glass thickness chamber slide with collagen. To avoid the meniscus which inevitably forms in small wells and precludes clear and focused live imaging, we designed and 3D-printed a “ collagen press” to flatten the collagen hydrogel as it sets. The press rests on the internal walls of the slide with glass-coated supports that hang 0.3mm from the well bottom (Fig 3B, Supplemental File 1), creating a uniformly flat hydrogel surface upon removal that is ideal for high-resolution time-lapse imaging of ISC cell cycle dynamics over 48 hours (Fig 3C, Supplemental Video).

ISCs were thinly seeded onto the collagen hydrogels to allow sufficient space to expand, then imaged starting 24 hours post-plating to mitigate the impact of potential contact inhibition. After 48 h live imaging by confocal microscopy, the fluorescence profiles of individual nuclei were tracked at each 10 min frame interval, with changes in mVenus and mCherry levels used to define lengths of each cell cycle phase (Figure 3D, Supplemental Files 2-4). Cell cycle phase lengths were measured in cells which completed a full cycle, measuring from the initial division of the mother cell through to subsequent division of each daughter cell and discounting cells that remained green for 20+ hours as this is consistent with cell cycle exit. We tracked 196 individual PIP-FUCCI cells and 264 SOX9KO-PIP-FUCCI cells to quantify SOX-dependent lengths for each cell cycle phase. SOX9KO cells had significantly shorter median lengths for all cell cycle phases (41.3% shorter G1 phase, 13.3% shorter S phase, 12.7% shorter G2/M phase) culminating in a 21.1% decrease in median total cell cycle length (Fig 3E,F). This dynamic live imaging approach goes beyond the snap-shot approach of widely used DNA content analysis to quantitatively demonstrate that SOX9 negatively regulates cell cycle progression by significantly increasing all cell cycle phase lengths, with preferential lengthening seen for G1.

### SOX9 promotes stem cell activity in human ISCs

Since SOX9 is dynamically expressed throughout the stem and progenitor cell populations in the mouse and human crypt, we next wanted to test how SOX9 affects stem cell activity within human ISC monolayers. Organoid formation assays were used to test ISC function. For these assays, actively growing monolayers with different levels of SOX9 expression were dissociated to single cells then plated into Matrigel for eight days (Fig 4A). ISC function was determined by the ability of a cell to generate an organoid with a lumen. To determine whether higher proliferation correlates with higher stem cell activity across SOX9 levels, we tested SOX9KO, SOX9OE no Dox, and Sox9OE+Dox for ISC formation. Also, as we hypothesized that constant high SOX9 might preclude proliferation and organoid growth, a SOX9OE + Dox Pulse condition was also tested where cells received Dox prior to dissociation but none during organoid growth in Matrigel. Notably fewer organoids formed in SOX9KO ISCs compared to SOX9OE no Dox (Fig 4B,C). Cells expressing high SOX9 (SOX9OE + Constant Dox) survived through 8 days but failed to grow beyond small clumps. These were not quantified since their lack of growth made it difficult to discern live vs dead clusters of cells. SOX9OE + Dox Pulse cells formed the most organoids, indicating that high SOX9 promotes survival and stem cell capacity, but stopping the SOX9 overexpression is then necessary for the cells to re-enter the cell cycle and form healthy organoids. Organoid formation assays were quantified across n=7 wells for each condition in N=3 separate experiments (Fig 4C). Normalized to SOX9OE no Dox controls, SOX9KO wells formed 48.2% fewer organoids and SOX9OE + Dox Pulse wells formed 521% as many organoids. Thus, increasing organoid formation ability correlated with SOX9 expression at time of dissociation, consistent with a model wherein SOX9 promotes stem cell survival and self-renewal – often referred to as “ stemness”. The SOX9OE + Dox Rescue condition may mimic putative SOX9-high reserve stem cells, which express high SOX9 until injury (replicated by single cell dissociation), then dynamic regulation into lower SOX9 expression allows for proliferation to replenish the crypt populations. As such, the ability to act as a rISC may only be allowed through dynamic cell cycle regulation within intestinal crypts. Altogether, this assay indicates that high SOX9 increases stem cell ability in ISCs, while loss of SOX9 hampers ISC function.

**Figure 4:**
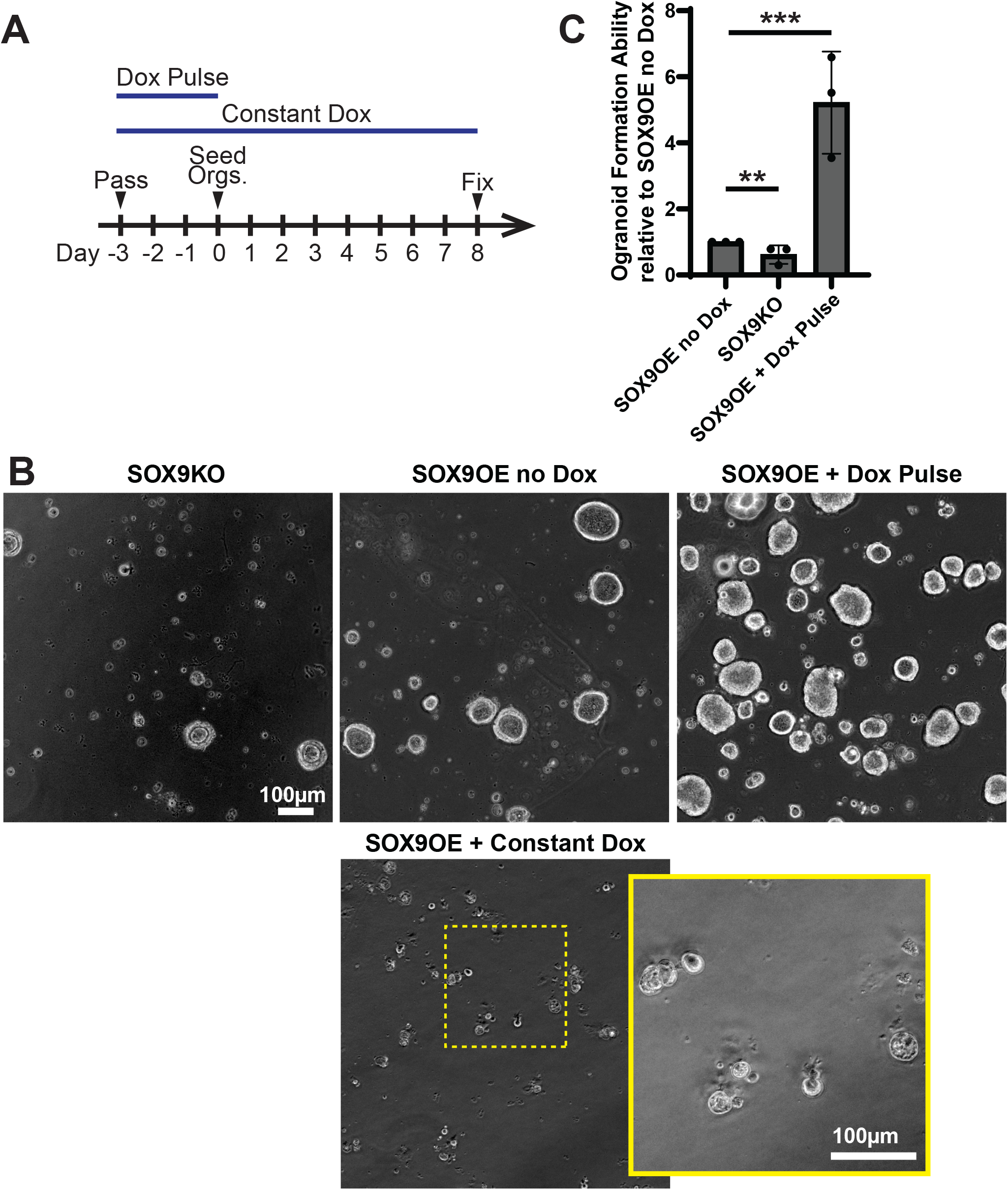
SOX9 promotes stem cell activity in human ISCs. **A)** Schematic timeline for organoid formation assays. Cells were passaged as monolayers, then Dox was added to the relevant conditions upon plating. Three days later, monolayers were dissociated to single cells and plated as organoids with/without Dox for 8 days prior to fixing and quantifying. **B)** Representative phase contrast images of organoids following 8 days of growth from single cells. Increased magnification inset shown for SOX9OE + Constant Dox organoids. All scale bars = 100 μm **C)** Bar graphs depicting average organoid formation ability across all conditions relative to SOX9OE no Dox. Statistics shown using T Test, P values: * < 0.05, ** <0.005, *** < 0.001.

### SOX9 represses human ISC differentiation into absorptive enterocytes and goblet, enteroendocrine, and BEST4^+^ cells, and promotes tuft cells and follicle-associated epithelium

We next tested how SOX9 expression affects the differentiation fate of human ISCs in vitro. WT, SOX9KO, Sox9OE, and SOX9KOOE monolayers were passaged into MM with/without Dox, then growth factors were removed for 5 days for cells to differentiate (Fig 5A). qPCR was performed for classical markers of major lineages to assay differentiation capacity for all intestinal cell types. Markers of the absorptive enterocyte lineage (*APOA4* and *FABP6*) were increased in ISCs with no SOX9, with low expression seen in ISCs with high SOX9 (Fig 5B). A similar pattern was seen for enteroendocrine cells (CHGA), goblet cells (*MUC2, SPINK4*), and BEST4^+^ cells (*BEST4*), with differentiation markers for each lineage highest in the SOX9KO condition (Fig 5C-E). Inversely, markers of tuft cells (*POU2F3, PTSG1*) and Follicle-Associated Epithelium (FAE; *TNFRSF11A, SOX8*), which include M Cells, increased with high SOX9 (Fig 5F,G). Paneth cell markers (*DEFA5, DEFA6*) were not strongly enriched at any condition (Fig5H). Although *DEFA5* and *DEFA6* are detected in SOX9OE conditions, only one sample from each biological triplicate showed expression, indicating that Paneth cells are not efficiently formed in our system. Interestingly, Lysozyme (*LYZ*) expression increased in all three of the triplicate samples for both SOX9OE conditions (Fig 5I). Since *DEFA5* and *DEFA*6 were only increased in one sample each, this strongly suggests that the *LYZ* expression is arising from non-Paneth cells. This is consistent with our findings from the healthy human intestine, where *LYZ* is expressed in FAE, BEST4^+^ cells, and tuft cells in the human intestine, making *LYZ* mRNA a poor indicator of Paneth cell presence for in vitro models of human intestinal epithelium^13^. Thus, it is likely that the *LYZ* observed in this system arises from FAE or tuft cells that are strongly induced in the SOX9OE conditions (Fig 5 F,G). Altogether, SOX9 expression was found to affect differentiation into all main intestinal epithelial lineages, repressing absorptive enterocytes, goblet cells, enteroendocrine cells, and BEST4^+^ cells, and promoting tuft cells and follicle-associated epithelium.

**Figure 5:**
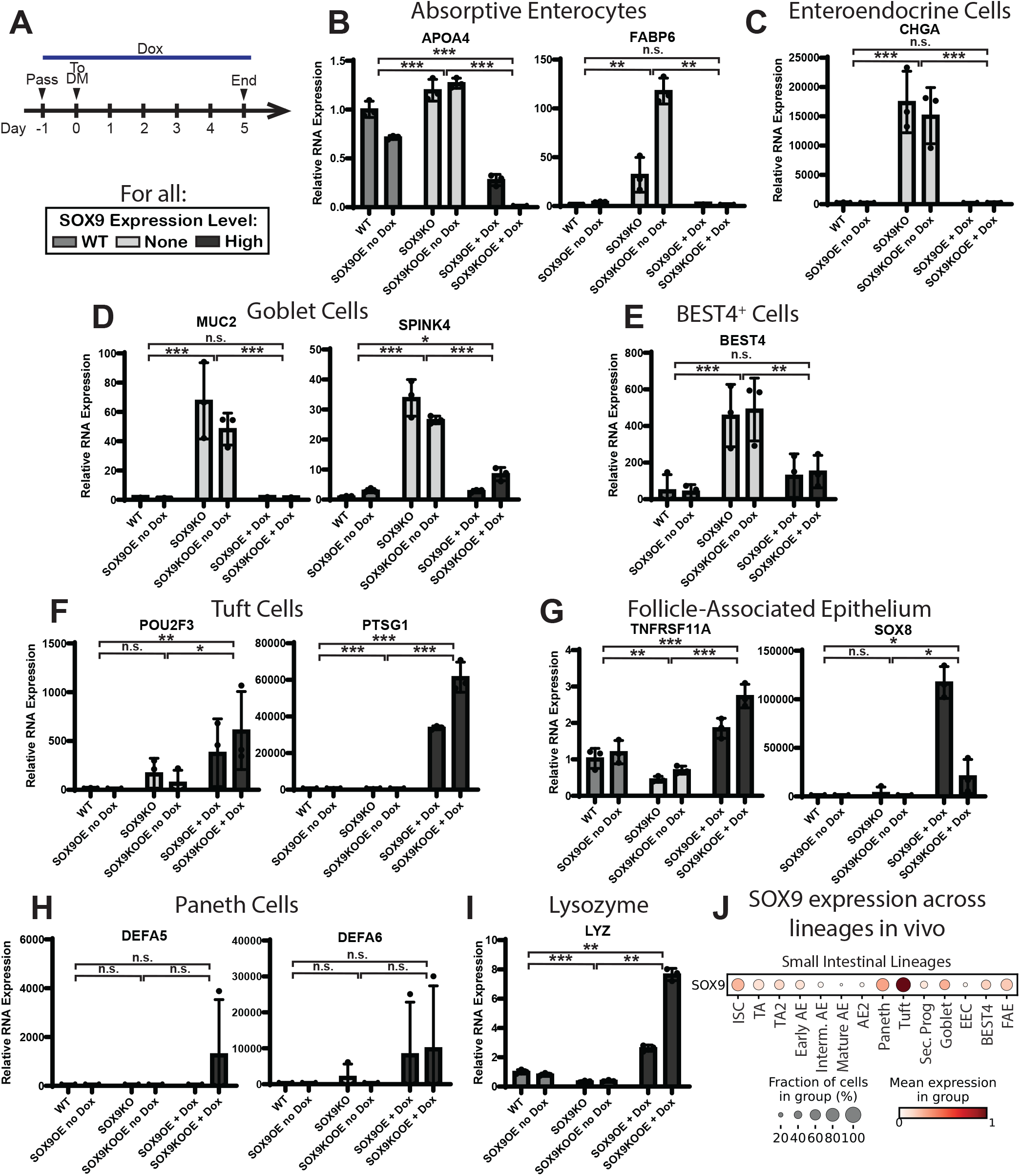
SOX9 regulates differentiation trajectory in human ISCs. **A)** Schematic timeline for differentiation assays. Monolayers were passaged, with Dox added to relevant conditions immediately upon plating in Maintenance Media. Cells were washed and cultured in Differentiation Media (DM) the following day and continuing for 5 days. **B-H)** Bar graphs depicting average transcriptional expression of classical markers of absorptive enterocytes (B), enteroendocrine cells (C), goblet cells (D), BEST4^+^ cells (E), tuft cells (F), follicle-associated epithelium (G), and Paneth cells (H). All bars are grouped by relative SOX9 expression: WT on left, KO in center, overexpression on right. **I)** Bar graph depicting average transcriptional expression of lysozyme across same SOX9 levels as B-H. For statistics for B-I, cell lines were grouped by functional SOX9 expression level: WT (WT and SOX9OE no Dox), KO (SOX9KO and SOX9KOOE no Dox), and OE (SOX9OE+Dox and SOX9KOOE+Dox). T Tests comparing each condition combined the triplicate values for each cell line within the condition into N=6 for the total condition. P values: * < 0.05, ** <0.005, *** < 0.001. **J)** Expression of *SOX9* across all small intestinal lineages from three healthy human donors^13^.

As a confirmation that our in vitro findings are physiologically relevant, we compared the above effects of SOX9 on differentiation to our recent atlas of the human gut epithelium^13^ (Fig 5J). *SOX9* expression in primary lineages from healthy intestines largely correlated with our in vitro results. In vivo data shows notable *SOX9* expression in tuft cells, FAE, Paneth cells, and ISCs, with levels dropping as the ISCs differentiate into TA or secretory progenitor cells. This is consistent with our in vitro results showing SOX9 promotes stemness and differentiation of tuft cells and FAE, with possible low-level promotion of Paneth cells. In vivo data also shows low *SOX9* expression in absorptive enterocytes and enteroendocrine cells, consistent with our in vitro results showing that SOX9 represses differentiation into these populations. Two points of interest are goblet cells and BEST4^+^ cells, which both express moderate levels of *SOX9* in vivo even though our differentiation assay indicates that SOX9 represses differentiation into both lineages. This may indicate that SOX9 functions in mature cells of these lineages distinct from its role in fate specification. This idea is supported by the low SOX9 seen in secretory progenitors which are believed to mature into tuft, Paneth, and goblet cells, all of which have higher levels of SOX9^13^. Altogether, this suggests that the effects of SOX9 on differentiation in our system are likely relevant to physiological differentiation as opposed to being artifacts of our in vitro system.

## DISCUSSION

The decision of an ISC to self-renew or differentiate is a complex genetic and molecular process that integrates extrinsic and intrinsic signals. The sum of these inputs can tip the decision toward an ISC continuing through another round of the cell cycle or exiting cell cycle to become a post-mitotic lineage. How an ISC regulates, interprets, and uses cell cycle phases to influence fate decisions is not understood. Here, we investigate SOX9-dependent effects on ISC proliferation at the resolution of cell cycle phases with the goal of laying the foundation for more granular mechanistic studies aimed at determining the genetic and epigenetic factors that underpin the decision to maintain a cycling state or exit to differentiation.

In the mouse intestine, loss of Sox9 has repeatedly been shown to increase the number of proliferative cells in the crypt^7, 22, 23, 36^, yet none of these studies analyzed the effects of Sox9 on cell cycle phase length. All available studies of SOX9 in any system currently rely either on measuring total proliferation or on DNA content analysis to analyze the effects of SOX9 on cell cycle in their respective systems. While DNA content analysis is a useful readout for cells within the cell cycle, it cannot accurately describe the effects of cells leaving the cell cycle. For example, if SOX9 increases cells within the G0/G1 population, this alone does not inform whether the G1 phase is lengthened in actively proliferating cells or whether more cells are exiting the cell cycle into quiescent G0 state but the lengths of all phases in actively cycling cells remain the same. Highly accurate Fluorescent Ubiquitination-based Cell Cycle Indicator (FUCCI) reporters, introduced in 2008, enable live imaging and quantification of each cell cycle phase. Several iterations of FUCCI reporters have been developed in the following years^35, 37^. Here, we used a PIP-FUCCI allele^35^ to quantify phase lengths in human ISCs and show that loss of SOX9 causes a predominant 41.5% shorter G1 along with reduction in length of the entire cell cycle and reduced lengths of S and G2/M phases.

G1 is a key phase of the cell cycle where ISC decisions revolving around undergoing another round of cell division or differentiating are made. Perhaps most notable is the Retinoblastoma (RB1) restriction point where the dephosphorylated RB1 tumor suppressor protein keeps a cell in G1 until cellular processes are in place for progression to S phase. In mice, Lgr5+ ISCs remain in an unlicensed G1 state prohibiting S phase entry, and it has been proposed that this elongated G1 allows cell fate decisions to be made^38^. Chromatin remodeling has also been shown to occur in G1^39^, and changes to transcription factor binding ability have even attributed to cell cycle proteins in G1^25^. We find that altered SOX9 expression levels regulate differentiation into every lineage in our in vitro model. Interestingly, cell lineages sort into biased lineages based on SOX9KO (fast G1) or SOX9OE (slow G1) environments. If SOX9-dependent G1 length is temporally regulating key G1 events that specify lineages, then absorptive enterocytes, goblet, and enteroendocrine cell specification events may favor shorter G1 lengths and tuft, Paneth, and FAE specification events may favor longer G1 lengths. Our study raises the intriguing possibility that SOX9 might regulate ISC differentiation indirectly through its broad effects on cell cycle, with longer/shorter G1 phase lengths priming cells to receive and respond to different differentiation factors, as has been shown to be the case in embryonic stem cell models^27-29^. While not explored in this study, uncoupling G1 lengths from SOX9 levels by pharmacological or genetic means would be necessary to test this hypothesis.

While we show that many SOX9-dependent differentiation effects are consistent with in vivo expression of SOX9 in the human intestine, we also find inconsistencies from mouse studies. Two independent studies evaluate lineage bias in a Sox9-flox mouse model^22, 23^. One shows Sox9-flox mice have fewer goblet cells while the other shows goblet cells unchanged, with both studies finding that enteroendocrine cell numbers are unaffected by Sox9 loss. This is inconsistent with our results showing drastically increased expression of goblet and enteroendocrine cell differentiation markers in SOX9KO monolayers. As we have shown numerous differences in mouse and human lineages in vivo^40^, species differences could contribute to these apparent inconsistencies. Alternatively, the differences could be related to extrinsic influences that occur during G1 in the mouse but are lacking in our human in vitro system. The precise timing of crucial changes in SOX9 expression as cells develop in vivo was not tested in our human ISC culture system but could also be essential for lineage specification. The flexibility and sensitivity of our human ISC platform will facilitate studies to test these possibilities.

One intriguing result from our studies is the drastic effect that SOX9 has on organoid formation ability in healthy human ISCs. SOX9KO cells formed significantly fewer organoids than those with WT levels of SOX9. Lack of stemness in SOX9KO cells is consistent with a study on mouse hair follicles, where loss of Sox9 decreased maintenance of quiescent stem cells, causing stem cells to have less regenerative capacity and deplete over time^41^. In our study, continuous induction of high SOX9 in human ISCs caused single cells to survive plating into 3D Matrigel cultures, but they did not proliferate appreciably until after removal of doxycycline allowed reduction of SOX9 levels. Notably, after removal of Dox, the ISCs reinitiated cycling and displayed over 5-times increased organoid forming activity compared to control conditions. This shows that high SOX9 levels do not promote terminal cell cycle exit, but rather support a reversible non-proliferative state while simultaneously preserving stemness. Collectively, these are properties that define reserve ISCs (rISCs)^42-44^.

rISCs in the murine intestine are well studied^2, 5-11, 42^, yet few mechanisms are shown that confer their properties and function. Our group previously showed that loss of Sox9 in the mouse epithelium caused loss of rISC activity^7^. While the genetic ablation model was incapable of attributing the loss of rISC activity to Sox9-high cells, we demonstrated that there was a loss of slowly dividing label-retaining cells (LRCs), which are Sox9-high cells consistent with a secretory precursor that predominantly gives rise to Paneth cells and exhibits rISC activity post-irradiation damage^21^. We also showed that Sox9-high cells exhibited more co-expression with the general proliferation marker Ki67 following high dose IR in a Sox9-EGFP reporter mouse, suggesting this population was re-entering the cell cycle^20^. There was no increase in the number of Sox9high cells in these mice during post-irradiation regeneration, suggesting that Sox9 levels reduced in these cells prior to cell division. This is consistent with our SOX9OE + Dox Pulse organoid formation results, supporting the potential role of SOX9 as a regulator of rISC activity. A non-genetic link that points to the G1 phase as being protective for ISCs was demonstrated when the restriction point inhibitor, Palbociclib, was used to arrest ISCs in G1 in mice^45^, causing substantial retention of ISC activity and recovery post-irradiation. This suggests that ISCs in G1 have increased rISC potential and supports the interpretation that SOX9-dependent rISC activity is conferred by elongation of G1. While linking high SOX9 levels to rISC activity will require future irradiation studies on our human ISC lines, our current findings are highly translational as related damage response mechanisms involving SOX9 have been noted across the gastrointestinal tract^42, 46, 47^. These findings thus lay the groundwork for further studies analyzing this mechanism to determine factors that regulate the stem cell injury response throughout the intestine and beyond.

## METHODS AND MATERIALS

### Tissue procurement, dissection, and crypt isolation

Intestinal epithelial cells were harvested from human intestines following a published protocol^33^. Briefly, donor-grade human intestines were received from HonorBridge (formerly Carolina Donor Services). Jejunum was defined as the upper half of the small intestine once the first 9cm (duodenum) was removed. A 3×3 cm piece was resected from the middle of the jejunal length, then stored in Advanced DMEM/F12 + 10 μM Y27632 and 200 μg/mL Primocin on ice until crypt isolation. To reduce mucus, the tissue was incubated in PBS + 10 mM N-acetylcysteine for 15 min then transferred to Isolation Buffer (5.6 mM Na2HPO4, 8.0 mM KH2PO4, 96.2 mM NaCl, 1.6 mM KCl, 43.4 mM Sucrose, and 54.9mM d-sorbitol) + 2 mM EDTA + 0.5mM DTT for 30min with gentle rocking at room temperature then vigorous shaking for 2 min. After shaking, the tissue was transferred to a new tube of Isolation Buffer + EDTA + DTT and the rocking then shaking were repeated. This was repeated six times, with supernatant for each stored on ice. Following the shakes, supernatants from each round were checked via light microscope for the presence of crypts and villi. Crypt-enriched shakes were pooled, washed in Isolation Buffer, then cultured or frozen.

### Tissue Culture

Human IECs were cultured on collagen plates prepared following a published protocol^31^. Maintenance Media was prepared as previously defined^30^. Briefly, L cells expressing transgenic Wnt3A, Noggin, and R-spondin3 (ATCC CRL-3276) were cultured in Collection Media (20% Tetracycline-negative Fetal Bovine Serum, 1% Glutamax, 1%Pen/Strep, in Adv DMEM/F12) for 12 days, with media collected daily. Maintenance Media consisted of 50% conditioned Collection Media and a final concentration of 2% B-27 Supplement, 5 mM Nicotinamide, 10 mM HEPES, 1 mM Glutamax, 1X Pen/Strep, 0.6125 mM N-Acetylcysteine, 25 μg/mL Primocin, 1.5 μM, 25 ng/mL mEGF, 1 nM Gastrin, and 5 nM Prostoglandin E2. Fresh crypts were plated with 200 mg/mL Primocin, 200 mg/mL Gentamycin, and 0.5 mg/mL amphotericin B for the first week. Cells were grown in a humidified incubator at 37 °C with 5% CO_2_.

Cells were passaged every 4-7 days, generally at ratios of 1:3 or 1:4. For passaging cells from a six-well collagen plate, the collagen patty and 1 mL of media from each well were incubated in a 37 °C water bath with 100 μL of 5,000 U/mL collagenase IV until collagen completely dissolved, centrifuged at 800 xG for 2 min, washed in dPBS, then digested in 1 mL TrypLE Express + 10uM Y27632 for 5 min in a 37 °C water bath. Cells were dissociated by triturating with a P1000 pipet tip then quenched with FBS. Cells were then resuspended in Maintenance Media + 10uM Y27632 and replated.

For differentiation fate analysis, cells were passaged into MM including 100ng/mL Dox if relevant. The next day, cells were washed three times in warm AdvDMEM/F12 then plated in Differentiation Media (DM) with or without Dox for 5 days prior to RNA harvest (see below). DM consisted of a final concentration of 10 mM HEPES, 2 mM Glutamax, 1x Pen/Strep, 1.25 mM N-Acetylcysteine, 50 μg/mL Primocin, 50 ng/mL mEGF, and 500uM A83-01 in AdvDMEM/F12. SOX9OE monolayers were treated with 100ng/mL doxycycline for at least 48h to induce SOX9 overexpression.

### Genetic Engineering

All genetic engineering was completed following previously described protocols^33^. Briefly, DNA segments of interest were isolated using restriction enzymes or amplified using CloneAmp HiFi PCR Premix. Plasmids were generated using an In-Fusion HD Cloning Kit then isolated from bacterial stocks using a QIAGEN HiSpeed Maxi kit. pPIGA-PHD was a gift from Linzhao Cheng (Addgene plasmid # 26778 ; http://n2t.net/addgene:26778; RRID:Addgene_26778)^48^. sg resistant gamma-tubulin was a gift from Maria Alvarado-Kristensson (Addgene plasmid # 104433 ; http://n2t.net/addgene:104433; RRID:Addgene_104433)^49^

For CRISPR/Cas9 transfections, crRNA and tracrRNA oligos were purchased from IDT and resuspended at 100 pmol/mL. 2 μL (200 pmol) of each of gRNA and tracRNA were combined with 2 μL of 5X annealing buffer and 4 μL nuclease-free dH2O and annealed in a thermocycler (95 °C – 5 min, 95°C to78°C at -2 °C/s, 78 °C – 10 min, 78 °C to 25 °C at -0.1 °C /s, 25 °C – 5 min), then transferred to ice. 3 μL (60 pmol) of this mix was added to 2 μL TrueCut Cas9v2 (approximately 60 pmol), incubated at room temperature for 15 min, then stored on ice until use.

Plasmids were electroporated into human cells using the Neon Transfection System with a 100 μL Kit. Cells were resuspended in 100 μL Neon Buffer R at 10,000 cells/μL with 6 μg of the plasmid of interest. For PIP-FUCCI and SOX9OE constructs, Super PiggyBac Transposase Expression Vector was added at 5ng/μL. For SOX9KO, cells were resuspended with 60 pmol Cas9 + 60 pmol annealed crRNA:trac:RNA. Cells were electroporated using Neon preset #5 (1,700 V, 1 pulse, 20 ms) then immediately added to a six-well collagen plate with 3 mL Maintenance Media + 1:1000 Y27632. Around 4-7 days post-transfection, colonies were selected using their respective antibiotics (Blasticidin 10 μg/mL, G418 200 μg/mL, hygromycin 200 μg/mL) for 4-8 days. Clones were then isolated by digesting the collagen patty with 5,000 U/mL collagenase IV at 100 μL/mL at 37 °C for 25 min. Colonies were washed with dPBS then individual colonies were picked using a 20 μL pipet over a light microscope and placed into individual wells of a 48-well collagen plate with 300 μL Maintenance Media + 1:1000 Y27632.

### Immunostaining

Fresh human intestinal tissue was fixed in 4% paraformaldehyde (PFA) overnight at 4 °C then transferred to 70% ethanol the next day. Tissues underwent sectioning and placement onto glass slides for downstream analysis. For staining, sections underwent routine deparaffinization and rehydration using Histoclear then an ethanol gradient. Slides were then washed with running water, permeabilized with 0.3% Triton X-100, blocked using 3% BSA in PBS, then primary antibodies were added in 3% BSA overnight at 4 °C. The following day, slides were washed 3x with PBS, secondary antibody and bisbenzimide were added in 3% BSA for 1 h at room temperature, then slides were washed in plenty of PBS and mounted using ProLong Gold antifade reagent. Sections were imaged on a Zeiss LSM900 inverted confocal microscope using a 20X/0.80 Plan Apo objective at 405, 488, and 640nm excitation.

Stem cell monolayers were fixed for 20 min with 4% PFA at room temperature then washed in dPBS and stored at 4 °C until staining. For staining, cells were permeabilized with 0.5% Triton X-100, washed with 0.75% glycine in PBS to quench free formaldehyde, blocked using 3% BSA in PBS, then primary antibodies were added in 3% BSA overnight at 4 °C. The following day, cells were washed 3x with Wash Solution (0.1% BSA, 0.2% Triton X-100, 0.05% Tween-20 in PBS), secondary antibody and bisbenzimide were added in 3% BSA for 1 h at room temperature, then cells were washed in plenty of PBS and stored at 4 °C in the dark until imaging. Plated cells with immunofluorescence staining were imaged on a Keyence BZ-X800 fluorescent microscope using a LD 20x/0.45 Plan Fluorite Keyence objective and 49000-ET-DAPI. 49002-ET-GFP, or 49006-UF1-ET-CY5 Filter cubes from Chroma. When multiple images are shown in a figure panel, all were taken with the same microscope settings and had the same display adjustments made. Images were analyzed with BZ-X800 analyzer software or FIJI software^50^.

### RNA Extraction and qPCR

For RNA extraction, monolayers were treated with collagenase IV at 100 μL/mL at 37 °C for ∼25 min. Cells were then pelleted, washed with dPBS, pelleted again, then resuspended in 300 μL lysis buffer from an RNAqueous Micro Total RNA Isolation Kit. RNA was isolated using the kit following the manufacturer’ s instructions. Concentration of the eluted RNA was measured on a Qubit 3 Fluorometer using a Qubit™ RNA High Sensitivity (HS) Assay Kit. cDNA was made from 50ng RNA using iScript™ Reverse Transcription Supermix, following manufacturer’ s instructions.

Gene expression levels were assayed via quantitative polymerase chain reaction (qPCR) on a StepOnePlus™ Real-Time PCR System or using Fluidigm BioMark HD high throughput qPCR using Flex Six Gene Expression IFC chips. Both systems used Taqman Probes (SOX9 Hs00165814_m1, LYZ Hs06596823_s1, FABP6 Hs01031183_m1, APOA4 Hs00166636_m1, CHGA Hs00900373_m1, BEST4 Hs00396114_m1, MUC2 Hs03005088_m1, SPINK4 Hs00205508_m1, DEFA5 HS00360716_m1, DEFA6 Hs00427001_m1, POU2F3 Hs00205009_m1, PTSG1 Hs00377726_m1, TNFSRF11a Hs00921372_m1, SOX8 Hs00232723_m1).

### Single cell dissociation and Flow Cytometry

To dissociate monolayers to single cells, collagen patties were digested and cells washed as described above. Following the 5 min digestion in TrypLE at 37 °C, cells were triturated with a P1000 pipet 20x, drawn up and forcibly expelled from a 28G syringe 3x to dissociate to single cells, then quenched with 10% FBS in Advanced DMEM/F12. Single cells were passed through a 30 μm filter, quantified on a hemacytometer with Trypan Blue, if relevant, then washed and pelleted.

To quantify 5-ethynyl-2’ -deoxyuridine (EdU) uptake using flow cytometry, growing cells were pulsed with 10 μM EdU for 1 h prior to dissociation, then fixed in 10 mL ice cold 4% PFA for 20 min at 4 °C with heavy rocking. Fixed cells were then washed and stored in Advanced DMEM/F12 at 4 °C. Following permeabilization, cells were stained for EdU using EdU Reaction Buffer (4 mM CuSO4, 2 μM Sulfo-CY5-azide, 0.2 M Ascorbic Acid, in PBS) for one hour at room temperature protected from light. Cells were then washed in PBS then sorted.

For DNA content analysis, cells were fixed in 10 mL ice cold 70% ethanol for 30min at 4 °C with heavy rocking. Fixed cells were washed and stored in Advanced DMEM/F12 at 4 °C. Cells were treated with 10 μg/mL ribonuclease at 37 °C for 20 min, washed in Advanced DMEM/F12, then 6 μg/mL propidium iodide was added 10 min prior to flow cytometry. All flow cytometry was run on a Sony SH800ZF cell sorter. For cell cycle analysis using propidium iodide, damaged cells were gated against using off-target fluorescence from the unused APC channel, then propidium iodide levels in single cells were used to calculate cell cycle via the Dean-Jett-Fox Model on FlowJo v10.7.1.

### Organoid Formation Efficiency Assays

Cells were treated with or without 100 ng/mL Dox for three days as monolayers. Monolayers were then dissociated to single cells following the same protocol as for flow cytometry above. Live single cells were quantified on a hemacytometer with Trypan Blue, then cells were pelleted and resuspended in Matrigel. 3000 live single cells were plated in one 10 μL Matrigel droplet per well on a 48-well culture plate. Droplets were inverted and allowed to solidify at 37 °C for 20 minutes, then 250 μL maintenance media was added with or without 100 ng/mL Dox. Organoids were grown for eight days with media changed every three days. On the eighth day, media was replaced with warm 4% paraformaldehyde for 25 min to fix the organoids, then plates were scanned on a Keyence Imager using bright field microscopy on a 4x objective. Organoids were counted by hand using the Cell Counter feature on Fiji software^50^. Organoids were only counted if they grew large enough to form a lumen. Seven wells were counted for each condition across three separate experiments (21 total wells per condition). Data is shown as the average count for each condition across experiments relative to levels in SO9OE + no Dox controls. Statistical differences were calculated using two-tailed T tests across all 21 replicate wells for each condition.

### Fluorescence Live Imaging and Analysis

Freely-growing monolayers stably transfected with the PIP-FUCCI fluorescent reporter were plated and imaged live on μ-Slide 4 Well ibiTreat chamber slides. For optimal imaging, slides were coated with an even 0.3 mm layer of collagen. To make these, a “ collagen press” was designed to sit on the well walls with ‘ feet’ hanging into the wells, leaving a 0.3 mm space for collagen. This press was 3D-printed using acrylonitrile butadiene styrene on a Stratasys F170 FDM (Fused Deposition Modeler) (Supplemental File 1). The feet were covered with coverslip glass using Glass Glue then coated with Rain-X following the manufacturer’ s instructions. Collagen was prepared as described in^32^. 300 μL collagen was added to each chamber, the press was set on top (Fig 3B), then the chamber slides with press were transferred to a 37 °C incubator for 1 h to set. The presses were then carefully removed and dPBS was added to cover the collagen hydrogel in each chamber. Coated slides were stored at room temperature in an airtight zip-top bag until use. Cells were passaged into the chambers one day prior to imaging, with media changed the following morning before imaging.

Live imaging was performed on an Andor Dragonfly Spinning Disk Confocal Microscope mounted on a Leica DMi8 microscope stand, using a Leica HC Fluotar L 25X/0.95 W 0.17 VISIR water objective. The pinhole size was set to 40 μm. The camera was an Andor iXon Life 888 EM-CCD, with Electron Multiplying Gain set to 150, Horizontal Shift Speed of 10 MHz – 16 bit, 2X Pre Amp Gain, 2.2 μs Vertical Shift Speed, Normal Vertical Clock Voltage and binned 2×2. For mCherry imaging, a 561 nm laser was used for excitation, and light was collected with a 593/43 Semrock emission filter. For mVenus imaging, a 514 nm laser was used for excitation, and light was collected with a 538/20 Semrock emission filter. Images had 512 × 512 pixels and the size of each pixel was 1.00 μm. At each position, Z stacks were acquired using a piezo Z stage with 5 μm intervals spanning 60 μm (13 steps). 2×2 montages were acquired at each position. Each Z stack in the 2×2 montage overlapped its neighboring Z stacks by 10%. Stitching was performed automatically in Andor’ s Fusion software (version 2.3.0.54), based on the signal from the YFP channel, using the High-Quality settings with both Background Subtraction and Auto Filter Width on. Four locations with a 2×2 montage Z stack in each were acquired per experiment. 2×2 montage Z stacks at each location were acquired every 10 minutes for 48 hours. All images directly compared to each other were acquired using the same settings. Typical settings were 2 s exposures with 10% laser power for mCherry, and 200 ms exposures with 0.5% laser power for mVenus. Temperature was maintained at 37 °C with an Okolab microscope enclosure, with continuous monitoring and feedback. 5% CO2 was warmed to 37 °C in the enclosure and humidified before being delivered to an enclosed stage top holder that contained the sample. To minimize evaporation during the experiment, 8 caps from 15mL falcon tubes were filled with water and placed surrounding the sample inside the stage-top sample holder. To maintain the water column on the objective, a Leica Water Immersion Micro Dispenser was used. Water dispensing settings were tested in mock experiments to ensure no break in the water column would occur during a 48-hour experiment. At the beginning of an experiment, water was delivered with the Micro Dispenser to create a large drop that fully covered the top of the objective. The Micro Dispenser reservoir was filled and connected to an additional beaker with distilled water using a communicating vessels design that fed the reservoir from the beaker as water levels dropped in the former. The Micro Dispenser was controlled by specialized Leica software throughout the experiment and programmed to deliver a 1s 100V pulse every 1 min 15s. Water levels in the reservoir and additional beaker were checked and refilled approximately every 12 hours.

Stitched Z stacks were analyzed using Bitplane Imaris software (version 9.9.1) and Microsoft Excel. A full analysis protocol, a custom Excel analysis workbook, and a sample analyzed Imaris file are included as Supplemental Files 2-4. First, Z slices without nuclear fluorescent signal were cropped out, and all remaining z stacks were max projected along the XY plane. Second, individual nuclei at each timepoint were marked manually using the “ spots” function in Imaris. Third, tracks were generated to connect the positions of nuclei marked with spots throughout the experiment. Fourth, mean fluorescent intensities were exported from Imaris into a custom Excel workbook that simplified determination of cell cycle phases. Cells were only tracked from mother cells which divided at least 5 h after the start of imaging to allow cells to acclimate to the imaging environment. Tracking was also only started on mother cell divisions that occurred at least 20 h prior to the end of the video. Cells were only tracked if they stayed in frame and in focus throughout their entire cell cycle, and cells were chosen from a variety of spots across the viewing area and the timelapse for tracking. An average of 90 nuclei were tracked at each location in an experiment (range=59-106 nuclei). The Excel workbook presents graphs of normalized mVenus and mCherry fluorescence. The G1-S transition was defined as the point when mVenus signal dropped over 50%. The S-G2 transition was defined as the point when the mVenus signal began rising above the lowest maintained level again. Final graphs and significance analyses were made using GraphPad Prism 9. Data is shown as violin plots for each cell cycle phase and for total cell cycle duration, with median values shown and Mann-Whitney testing used to define significance between pairs.

### scRNAseq analysis

Transcriptional signatures of small intestinal lineages were used from a scRNA-seq database containing 12,590 single cells from small intestine and colon across three patients^40^ to visualize gene expression across cell types. (Gene Expression Omnibus GSE185224).

## FIGURE LEGENDS

**Supplemental Video – Representative PIP-FUCCI ISC monolayer growth.** Representative 48 h video of a monolayer with PIP-FUCCI stably introduced into otherwise WT human ISCs. Each frame = 10 min, progressing at a rate of 4 hours per second.

**Supplemental File 1 – Design file for collagen press**

**Supplemental File 2 – PIP-FUCCI Analysis Protocol**

**Supplemental File 3 – Empty Analysis Workbook for PIP-FUCCI**

**Supplemental File 4 – Example PIP-FUCCI Analysis**

**Supplemental Table – Reagents and Materials**

## ACKNOWLEDGEMENTS

The author first and foremost thank the human donor and their family for the gift of tissue. The authors thank Steven Emanual through the Biomedical Engineering Department at UNC Chapel Hill for help with 3D printing the collagen presses. We thank Jean Cook and Jeremy Purvis for supplying the PIP-FUCCI construct and help with analyzing cell cycle. The authors thank Gabrielle Cannon and the UNC Advanced Analytics Core (UNC Center for GI Biology and Disease, P30 DK034987) for use of the FACS machine and their assistance with the bulk qPCR analysis in this study. The Microscopy Services Laboratory, Department of Pathology and Laboratory Medicine, is supported in part by the P30 CA016086 Cancer Center Core Support Grant to the UNC Lineberger Comprehensive Cancer Center. The Andor Dragonfly microscope was funded with support from National Institutes of Health grant S10OD030223.

This research was supported by a CGIBD pilot grant through funding from the National Institutes of Health, P30 DK034987, T32DK07737, and F32DK124929 (J.B.), T32GM133364 (K.A.B.), F30DK126307 (M.T.O.), R01DK115806 and R01DK109559 (S.T.M.) and the Katherine E. Bullard Charitable Trust for Gastrointestinal Stem Cell and Regenerative Research.

