## Supplemental Table for "SOX9 elongates cell cycle phases and biases fate decisions in human intestinal stem cells"

| Reagent | Company | Catalog Number |
|---|---|---|
| **Chemicals, Peptides, and Recombinant Proteins** | | |
| N-Acetylcholine (NAC) | Sigma-Aldrich | A9165 |
| Dulbecco's Phosphate Buffered Saline (dPBS) | Gibco | 14190-144 |
| Na2HPO4 | Sigma | S7907 |
| KH2PO4 | Sigma | P5655 |
| NaCl | Sigma | S5886 |
| KCl | Sigma | P5405 |
| Sucrose | Fisher BP | 220-1 |
| d-sorbitol | Fisher BP | 439-500 |
| Y27632 | Selleck Chemical | S6390 |
| Ethylenediaminetetraacetic acid (EDTA) | Corning | 46-034-CI |
| Dithiothreitol (DTT) | Fisher Scientific BP | 172-5 |
| Protease VIII | Sigma | P5380 |
| Advanced DMEM/F12 | Gibco | 12634-010 |
| Bovine Serum Albumin | Fisher Scientific | BP1600-1 |
| Fetal Bovine Serum Tetracycline Negative | Gemini | 100-800 |
| Primocin | Invivogen | ant-pm-05 |
| Gentamycin | Sigma-Aldrich | G1914 |
| Amphotericin B | Sigma-Aldrich | A2942 |
| Collagenase IV | ThermoFisher | LS004189 |
| TrypLE Express | Gibco | 12605-010 |
| GlutaMAX | ThermoFisher | 35050061 |
| HEPES | Corning | 25-060-CI |
| Primocin | Invivogen (VWR) | mspp-ant-pm2 |
| Pen/Strep | ThermoFisher | 15070063 |
| N-acetylcysteine | Sigma-Aldrich | A9165 |
| EGF, murine | Peprotech | 315-09 |
| Nicotinamide | Sigma-Aldrich | N0636 |
| B27 Supplement | ThermoFisher | 12587001 |
| Gastrin | Sigma-Aldrich | G9145 |
| Prostaglandin E2 | Peprotech | 3632464 |
| A 83-01 | Sigma-Aldrich | SML0788 |
| SB202190 | Peprotech | 1523072 |
| Doxycycline | Sigma-Aldrich | D9891 |
| 5-ethynyl-2'-deoxyuridine (EdU) | Molecular Probes | C10634 |
| Propridium Iodide | Invitrogen | P3566 |
| CloneAmp HiFi PCR Premix | Takara | 639298 |
| In-Fusion HD Cloning Kit | Takara | 638920 |
| QIAGEN HiSpeed Maxi kit | Qiagen | 12662 |
| Super PiggyBac Transposase Expression Vector | System Biosciences | PB210PA-1 |
| nuclease-free dH2O | Corning | 46-000-CI |
| 5X annealing buffer | ThermoFisher Scientific | 100061876 |
| TrueCut Cas9v2 | ThermoFisher Scientific | A36498 |
| blasticidin | Gibco | A1113903 |
| hygromycin | Invivogen | ant-hg-1 |
| G418 (Geneticin) | Invivogen | ant-gn-1 |
| Paraformaldehyde | Thermo Scientific | AC416780250 |
| Histoclear | National Diagnostics | HS-200 |
| Triton X-100 | Fisher Scientific | BP151-100 |
| Tween-20 | Bio-Rad | 1706531 |
| ProLong Gold antifade reagent | Invitrogen | P36930 |
| Glycine | Fisher Scientific | BP381-5 |
| RNAqueous™-Micro Total RNA Isolation Kit | ThermoFisher Scientific | AM1931 |
| Qubit™ RNA High Sensitivity (HS) Assay Kit | Fisher Scientific | Q32852 |
| iScript™ Reverse Transcription Supermix | Bio-Rad | 1708841 |

| | | |
|---|---|---|
| Trypan Blue solution | Sigma | T8154 |
| Ribonuclease A from bovine pancreas | Millipore Sigma | R4642 |
| CuSO4 | Fisher Scientific | S25286 |
| Sulfo-CY5-Azide | Lumiprobe | A3330 |
| Ascorbic Acid | Fisher Scientific | AC352681000 |
| **Antibodies** | | |
| i-67 Monoclonal Antibody (SolA15), APC | Invitrogen | 17-5698-82 |
| Rabbit anti-SOX9 | Abcam | ab185230 |
| Bisbenzamide H 33258 Bioreagent | Sigma | B1155 |
| **Cells** | | |
| L-WRN cells | ATCC | CRL- 3276 |
| **Hardware** | | |
| Neon Transfection System | ThermoFisher | MPK5000 |
| Neon Transfection System 100 µL Kit | ThermoFisher | MPK10096 |
| Fluorescence Microscope | Keyence | BZ-X800 |
| Qubit 3 Fluorometer | Invitrogen | Q33216 |
| StepOnePlus™ Real-Time PCR System | ThermoFisher | 4376600 |
| Flex Six Gen Expression IFC | Standard Biotools | 100-6308 |
| 28G Insulin Syringe | BD | 329424 |
| Celltrics 30µm filter | Sysmex | 04-004-2326 |
| Sony SH800ZF cell sorter | Sony | SN 1500005 |
| µ-Slide 4 Well ibiTreat chamber slides | Ibidi | 80426 |
| Glass Glue | Loctite | |
| Rain-X Glass Treatment | Rain-X | 800002250 |
| Andor Dragonfly Spinning Disk Confocal microscope | Andor | |
| Leica HC Fluotar L 25X/0.95 W 0.17 VISIR water objective | Leica | 506375 |
| **Software** | | |
| Sony Cell Counter Software | Sony | |
| Adobe Illustrator | Adobe | |
| GraphPad Prism 9 | GraphPad | |
| Microsoft Office 365 Suite | Microsoft | |
| Bitplane Imaris software (version 9.9.1) | Oxford Instruments | |
| BZ-X800 Analyzer | Keyence | |
